# Rescuing impaired hippocampal-cortical interactions and spatial reorientation learning and memory during sleep in a mouse model of Alzheimer’s disease using hippocampal 40 Hz stimulation

**DOI:** 10.1101/2024.06.20.599921

**Authors:** Sarah D. Cushing, Shawn C. Moseley, Alina C. Stimmell, Christopher Schatschneider, Aaron A. Wilber

## Abstract

In preclinical Alzheimer’s disease (AD), spatial learning and memory is impaired. We reported similar impairments in 3xTg-AD mice on a virtual maze (VM) spatial-reorientation-task that requires using landmarks to navigate. Hippocampal (HPC)-cortical dysfunction during sleep (important for memory consolidation) is a potential mechanism for memory impairments in AD. We previously found deficits in HPC-cortical coordination during sleep coinciding with VM impairments the next day. Some forms of 40 Hz stimulation seem to clear AD pathology in mice, and improve functional connectivity in AD patients. Thus, we implanted a recording array targeting parietal cortex (PC) and HPC to assess HPC-PC coordination, and an optical fiber targeting HPC for 40 Hz or sham optogenetic stimulation in 3xTg/PV^cre^ mice. We assessed PC delta waves (DW) and HPC sharp wave ripples (SWRs). In sham mice, SWR-DW cross-correlations were reduced, similar to 3xTg-AD mice. In 40 Hz mice, this phase-locking was rescued, as was performance on the VM. However, rescued HPC-PC coupling no longer predicted performance as in NonTg animals. Instead, DWs and SWRs independently predicted performance in 40 Hz mice. Thus, 40 Hz stimulation of HPC rescued functional interactions in the HPC-PC network, and rescued impairments in spatial navigation, but did not rescue the correlation between HPC-PC coordination during sleep and learning and memory. Together this pattern of results could inform AD treatment timing by suggesting that despite applying 40 Hz stimulation before significant tau and amyloid aggregation, pathophysiological processes led to brain changes that were not fully reversed even though cognition was recovered.

**Significance Statement:** One of the earliest symptoms of Alzheimer’s disease (AD) is getting lost in space or experiencing deficits in spatial navigation, which involve navigation computations as well as learning and memory. We investigated cross brain region interactions supporting memory formation as a potential causative factor of impaired spatial learning and memory in AD. To assess this relationship between AD pathophysiology, brain changes, and behavioral alterations, we used a targeted approach for clearing amyloid beta and tau to rescue functional interactions in the brain. This research strongly connects brain activity patterns during sleep to tau and amyloid accumulation, and will aid in understanding the mechanisms underlying cognitive dysfunction in AD. Furthermore, the results offer insight for improving early identification and treatment strategies.

## Introduction

Alzheimer’s disease (AD) is the leading cause of dementia worldwide, resulting in devastating effects for patients, caregivers, and society (McDade and Bateman, 2017; Association, 2019). Spatial navigation deficits are an early symptom of AD, and may be a better differential diagnostic marker than classical memory deficits (Henderson et al., 1989; Weintraub et al., 2012; Lithfous et al., 2013; Allison et al., 2016; Tu et al., 2017; Coughlan et al., 2018). These spatial navigation deficits could be the result of deficits in memory consolidation and formation (Billings et al., 2005; Cayzac et al., 2015; Mably et al., 2017), memory retrieval (Harrison et al., 2016; Roy et al., 2016), navigation computations (Cañete et al., 2015; Davis et al., 2017), or a combination of memory and navigation dysfunction.

Memory formation and encoding is highly dependent on sleep (Marshall and Born, 2007; Diekelmann and Born, 2010; Kim et al., 2019; Klinzing et al., 2019), and AD patients demonstrate a variety of sleep disruptions such as increased insomnia, increased wakefulness, increased sleep fragmentation, and decreased slow wave sleep (Bliwise et al., 1989; Grace et al., 2000; Bonanni et al., 2005; Peter-Derex et al., 2015; Holth et al., 2017). Many rodent models of AD recapitulate sleep changes (Wisor et al., 2005; Zhang et al., 2005; Schneider et al., 2014; Sethi et al., 2015; Kent et al., 2018). These sleep disruptions could be directly impacting AD patients ability to form and retain memories, and studies have found that AD memory symptoms may be a consequence of forgetting from one day to the next (Billings et al., 2005).

Memory consolidation involves coordinated interactions between the hippocampus (HPC) and cortex, which are critical for learning (Wang and Morris, 2010; Schwindel and McNaughton, 2011; Maingret et al., 2016; Skelin et al., 2018). It is possible that changes in this network could be a contributing factor to memory impairments in AD, given the evidence of aberrations in HPC-cortical connectivity (Wang et al., 2006; Allen et al., 2007; Park et al., 2017; Stoiljkovic et al., 2018; Cushing et al., 2020). It has been theorized that the HPC creates an index or code reflecting the spatio-temporal context of events, and that this allows the HPC to link the various aspects of a memory across cortex. Replay of these patterns within hippocampus, within cortex or across hippocampus and cortex during sleep is thought to play a role in building and strengthening the connections underlying this code (Teyler and DiScenna, 1986). Other theories of memory formation, such as multiple trace theory or trace transformation theory, also recognize the importance of HPC-cortical interactions early in the process (Nadel et al., 2000; Winocur et al., 2010). Thus, nearly all current theories emphasize the importance of HPC-cortical interactions in memory formation and strengthening.

Memory replay is critical for memory consolidation, and occurs in both HPC at the cellular level, and parietal cortex (PC) at the cellular and modular level (Louie and Wilson, 2001; Foster and Wilson, 2006; Ji and Wilson, 2006; Wu and Foster, 2014; Wilber et al., 2017; Cushing et al., 2020; Findlay et al., 2021). This replay involves cells firing during sleep in similar patterns to how they fired during experience repeatedly in a compressed time frame. Sharp wave ripples (SWRs) in HPC are a marker of this memory reactivation event, and disruption of SWRs result in impairments in memory formation (Jadhav et al., 2012; Nokia et al., 2012; Buzsáki, 2015; van de Ven et al., 2016; Roux et al., 2017; Fernández-Ruiz et al., 2019a; Zhen et al., 2021). Similarly, in cortex, we see delta waves (DW) and thalamocortical spindles, both of which are important for memory consolidation, and disruption or a reduction of these events leads to impaired memory formation (Siapas and Wilson, 1998; Nishida and Walker, 2007; Clemens et al., 2011; Latchoumane et al., 2017; Helfrich et al., 2018; Jiang et al., 2019; Kim et al., 2019). Furthermore, previous studies have shown that the temporal coordination of all of these events is also critical for memory formation (Siapas and Wilson, 1998; Sirota et al., 2003; Clemens et al., 2011; Staresina et al., 2015; Latchoumane et al., 2017; Helfrich et al., 2018; Skelin et al., 2018; Jiang et al., 2019; Cushing et al., 2020). AD has been shown to have impacts on not only these individual events (such as a reduction in SWR generation), but also on the temporal coordination of these events (Buzsáki, 2015; Ciupek et al., 2015; Caccavano et al., 2020; Cushing et al., 2020; Zhen et al., 2021). Specifically, we previously identified HPC-cortical interactions during sleep that predicted performance on a spatial navigation task the following day. Furthermore, these interactions were reduced in 3xTg-AD mice, which corresponded with deficits in learning the spatial navigation task (Cushing et al., 2020). Additionally, we found that Aβ load in HPC correlated with markers of memory reactivation better than Aβ load in PC. Thus, we attempted to clear pathology in HPC to see if we could ameliorate deficits in HPC-PC interactions during sleep.

One emerging potential treatment for AD involves stimulation of 40 Hz brain rhythms to reduce both tau and Aβ (Canter et al., 2016; Iaccarino et al., 2016; Singer et al., 2018; Adaikkan et al., 2019; Adaikkan and Tsai, 2020; Garza et al., 2020; Figueiro and Leggett, 2021; Guan et al., 2022). In human patients, this treatment is showing some promising results, including ameliorating sleep impairments (Jones et al., 2019b; Suk et al., 2020; Cimenser et al., 2021). In animals, multisensory stimulation has resulted in improved performance on memory tasks as well as reduction in amyloid beta (Aβ) and tau (Iaccarino et al., 2016; Adaikkan et al., 2019; Martorell et al., 2019; Garza et al., 2020). Optogenetic stimulation has similarly resulted in clearance of Aβ (Iaccarino et al., 2016). However, some other papers have found that light-only 40 Hz stimulation does not result in a reduction of pathology (Soula et al., 2023; Yang and Lai, 2023).

We implemented an optogenetic stimulation approach for HPC to attempt to ameliorate the deficits in HPC-PC interactions during sleep we previously identified. After mice had completed a daily spatial navigation task, and slept for memory consolidation, they underwent an hour long HPC optogenetic stimulation session, with either 40 Hz stimulation or sham stimulation. This occurred daily for 2-3 months (i.e., for the duration of the experiment).

We found that 40 Hz stimulation rescued both spatial navigation deficits and HPC-PC interactions during sleep. However, unlike in NonTg control mice, these measures were decoupled in 40 Hz mice, with HPC-PC interactions not predicting future spatial navigation performance. Instead, individual markers of memory reactivation, specifically SWRs in HPC and DWs in PC, now predicted subsequent performance. Thus, 40 Hz stimulation may rescue some aspects of cognition and brain function, but not restore the brain to its original state.

## Methods

### Animals

6-month female 3xTg-AD/PV^cre^ mice (APPSwe, PS1M146V, and tauP301L) were used. Sample size (n=7 for 40 Hz group, n=6 for sham group) was determined via power analysis to ensure that the lowest numbers of animals needed to be used for the proposed experiments. Power analysis was based on previously published behavior data, which showed the greatest variability for the planned analyses, and was performed using G*power (Faul et al.; Cushing et al., 2020). At this age, mice are expressing intracellular Aβ and tau, but plaques and tangles are not yet present. We ran one more 40 Hz mouse than sham as one 40 Hz mouse did not have full IHC performed. One 40 Hz mouse was hemizygous for the PV^cre^ gene, all other mice were homozygous. We included the hemizygous mouse in our analyses as the results were consistent with the homozygous mice, suggesting the hemizygous PV^cre^ expression level is sufficient.

## Behavior

### Pretraining

Mice were moved to single housing and water deprived to no less than 80% of their starting weight. *Alternation training* then began where mice learned to shuttle back and forth along a linear track for a water reward that was delivered in an enclosed start-box (i.e., rewarded for each traversal out to the end of the track and back to the start box). At the end of the track was a black barrier in front (from the mouse’s perspective) of a black background. The starting position was moved to varying locations by sliding the entire track, while the barrier remained fixed, resulting in variation in track length. Starting positions were randomly selected, between 56-76 cm from the reward zone. When mice met asymptote criteria (number of runs did not vary by more than ±6 on 3 of 4 days, or ran 50 or more times down the track in a single session), a date was scheduled for implantation of stimulating electrodes and the recording array. Mice continued to run the task every other day leading up to surgery, including the day before surgery.

### Surgical Procedure

Two bipolar stimulating electrodes were implanted unilaterally, targeting the left medial forebrain bundle (mfb; 1.9 mm & 1.4 mm posterior to bregma, ±0.8 mm lateral, 4.8 mm below dura). A 16-tetrode recording array (Chang et al., 2013) was implanted, targeting the right PC and dorsal hippocampus (2.2 mm posterior to bregma, 2.0 mm lateral). 1 µl of AAV carrying channel rhodopsin II (UNC, AAV-EF1a-DIO-hChR2(H134R)-EYFP) was slowly injected by hand into left hippocampus and left for 5 mins before slowly removing the needle 26-gauge needle (Hamilton 80383, -2.0 mm posterior to bregma, 1.8 mm lateral). An optical fiber (Thor Labs, FT200EMT) was implanted targeting left hippocampus (−2.0 mm posterior to bregma, 1.8mm lateral). AAV expression was confirmed via histological analysis after the animal was perfused. Animals recovered for two weeks, during which tetrodes targeting PC were moved down 31 µm daily for the first three days, and then every other day, to prevent sticking. Tetrodes targeted to PC were not advanced beyond the lower border of PC, as determined based on inspection of the LFP, and also depth records. Tetrodes targeting HPC were turned down up to 124 µm daily for the first three days, and then 31 µm every other day, until tetrodes reached hippocampus, as indicated by depth records, as well as the characteristic HPC LFP and single units.

### Stimulation Parameters

After recovery, animals were placed in a custom 44 x 44 x 44 cm black box with a nose poke port (Med Associates) in the left corner. Electrical stimulation lasting 500 ms was administered manually to shape the mice to use the nose poke port. Nose pokes were registered by a custom MATLAB program that automatically delivered stimulation (Guerrero et al., 2023). Once mice had been trained to use the nose poke port, stimulation parameters were varied (171-201 Hz, 30-130 µA current, electrode wire combination) to identify the settings that produced the highest response. No attempt was made to balance settings across genotype. However, as we have found from our previous studies (Stimmell et al., 2019; Cushing et al., 2020) this naturally leads to a response rate is comparable across genotype (t_(11)_=1.092, p=0.298), suggesting that differences in reward strength were not likely to contribute to observed effects. The same comparison was made for the two brain stimulation parameters that were adjusted, frequency (t_(11)_=0.31, p=0.762) and current (t_(11)_=0.86, p=0.408), which also did not vary across genotype.

### Spatial Reorientation Training

After optimal stimulation settings were identified, mice completed a spatial reorientation task previously described (Rosenzweig et al., 2003; Stimmell et al., 2019; Cushing et al., 2020). Briefly, the *spatial orientation task* is identical to *alternation training,* except for the addition of an 8 cm long goal zone in a fixed location in the room (**Fig. 1B**). For this task, if the mouse pauses in the real or virtual (for the virtual maze version described below, **Fig. 1B** *Bottom Right*) goal zone for a sufficient period, brain stimulation reward is delivered. The required duration of the pause in the goal zone gradually increases as the animal achieves asymptote criterion at each phase (0.5-2.5 sec in 0.5 sec increments). The real (or virtual) box and track are moved after each trial (sliding track or virtual teleportation), so the goal zone remains at a constant position within the real (or virtual) room; however, the goal zone can be at a variety of distances from the start box (40–110 cm). The velocity profile during the approach to the reward zone (slowing) is used to assess performance. Slowing in the reward zone suggests the mouse knows its location (Rosenzweig et al., 2003; Stimmell et al., 2019). Asymptote criteria was ±15% correct trials over 3 out of 4 consecutive days. Most data reported here comes from the next (virtual) task for which mice run many more trials per daily recording session (242% increase). Spatial reorientation training took 11-31 days, with variation due to mice meeting asymptote criteria on delays at different rates.

**Figure 1.**
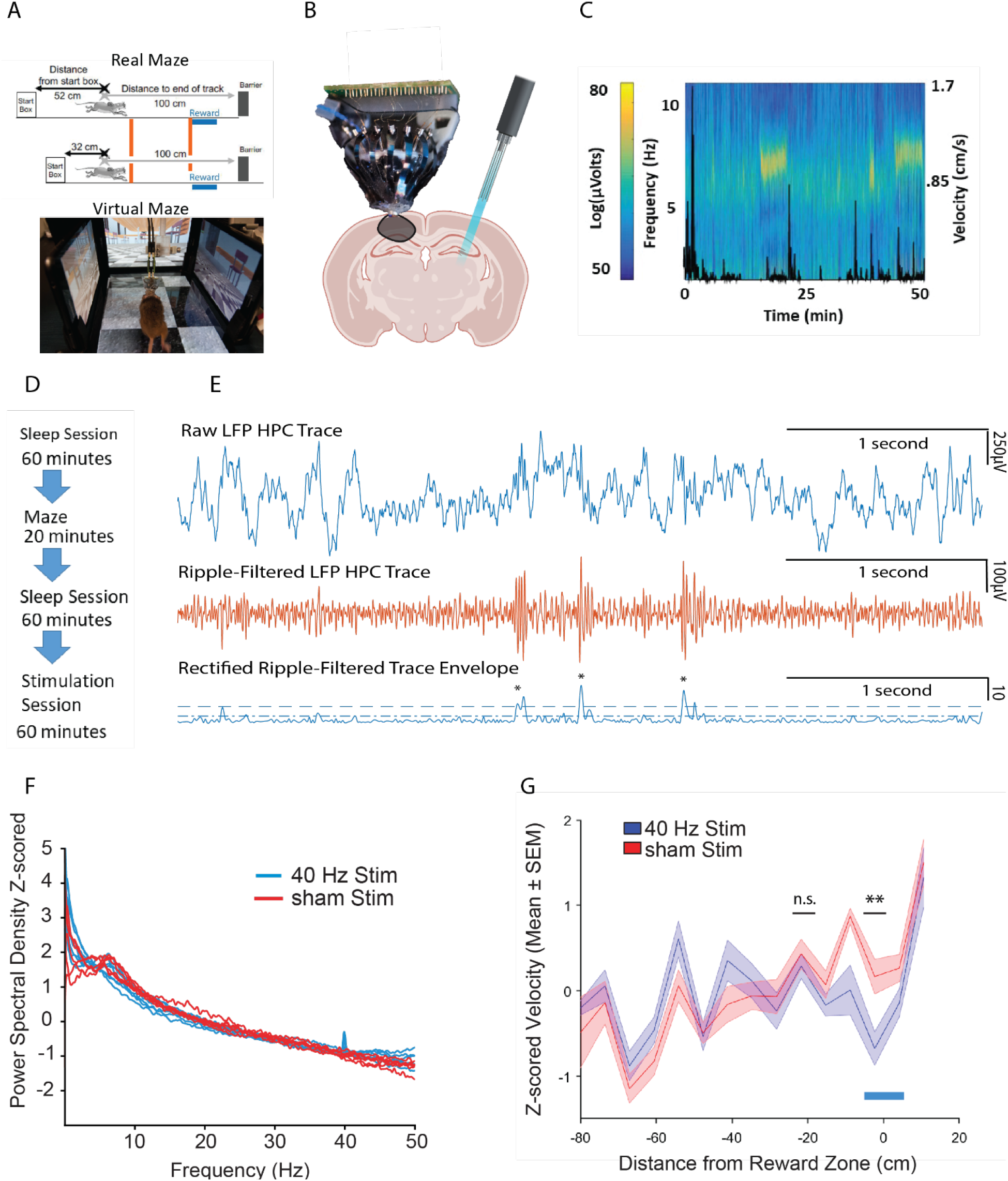
40 Hz stimulation of PV+ interneurons result in increases in 40 Hz power in HPC. **A.** The maze consists of a linear track with a start box and an unmarked reward zone (blue bar). Mice need to use external cues in the real or virtual room to locate the reward zone and slow to obtain a brain stimulation reward because local cues moved each trial (or were unavailable on the VM). Once a trial is completed, the start box is moved to a new random start location preventing self-motion alone from reliably indicating the reward zone. Adapted from Cushing et al., 2020). **B**. Schematic representing where stimulation and recording occurred. Optogenetic stimulation was targeted to the left hippocampus, and recording were obtained from the contralateral HPC and PC (circled). Created with BioRender.com **C.** Example time frequency power spectrum during sleep session used for sleep classification. Animal movement is indicated with the black line. Low power in theta band during stillness was considered SWS, high theta during still periods was considered REM.(Stan Leung, 1998; Lansink et al., 2008; Sirota et al., 2008; Mizuseki et al., 2009; Mizuseki et al., 2011; Grosmark et al., 2012; Cushing et al., 2020) Adapted from (Cushing et al., 2020). **D.** Each daily recording session consisted of 60 min pre-sleep session, followed by 20 min maze session, then a 60 min post-sleep session, and finished for a 60 min 40 Hz stimulation session. **E**. Example raw LFP trace from HPC (top) was filtered in the ripple band (75-300, middle), and an envelope was created with the z-scored filtered trace. Events with peaks >6 SD (dashed line) above the mean and 20-100ms were considered SWRs (Dragoi et al., 1999; Maier et al., 2003; Nimmrich et al., 2005; Jadhav et al., 2016; Oliva et al., 2016; van de Ven et al., 2016; Roux et al., 2017; Yamamoto and Tonegawa, 2017). Adapted from (Cushing et al., 2020). **F**. Representative example Z-scored power spectral density trace from HPC during the 1 hr long stimulation session for each 40 Hz mouse (blue) and each Sham mouse (red). **G**. Average z-scored velocity (mean ± SEM) as a function of distance from the reward zone (blue bar) for the first 6 days of the VM. Sham mice (red; n=6) slowed significantly less than 40 Hz stim mice (blue, p = 0.01; n=7).

### Virtual Maze

Upon completion of the spatial reorientation task, mice were trained on a virtual maze (VM) (https://www.interphaser.com/, Molina et al., 2016) as described above. The only difference was the addition of an acclimation period for learning to navigate the VM during head fixation. Note, prior to this point mice are head fixed daily for approximately 1 month while turning tetrodes, so acclimation to head fixation is already nearly complete. However, several mice did need additional time to acclimate to the virtual maze. Two 40 Hz stimulated mice needed an additional day to acclimate and only completed <4 trials on the first day, which was subsequently dropped from all analyses, and day 2 was counted as the start day. The acclimation period during VM training is to allow the animal to learn how to control the VM environment while head fixed.

Most mice prefer a specific head orientation for navigating the VM and it takes some time to find the optimal head orientation for each mouse. Head fixation was performed by hand fixation throughout the VM (Note, hand fixation means the fixation device was hand held. The actual mouse was not touched during hand fixation.) The tablet was coated with a thin layer of mineral oil and the mouse was allowed to run on the tablet in place. This is also how tetrode turning takes place, thus habituating the mouse to head fixation. The mouse’s paws were sensed on the floor tablet similar to a finger swipe on a cell phone screen, which caused the virtual environment to move at the same rate as if the mouse was running across a real floor. Virtual movement was restricted to forward or backward only (the virtual environment could not turn). Three wall tablets (front and both sides) were synced to the floor tablet to display the view of the virtual room. VM took 7-30 days, with variation again due to mice learning the task at different rates.

### Optogenetic Stimulation

At the end of each recording session (4-6 hours after light cycle began), mice remained in the sleep box for 60-70 min while optogenetic stimulation, either 40 Hz or sham, was performed (**Fig. 1B** contralateral hemisphere from recording array). Sham mice received a random poison shaped distribution of pulse spacings designed so that the average pulse spacing was 40 Hz. They received the same number of pulses, but with no consistent frequency entrainment as reflected in the lack of 40 Hz power in these mice (**Fig. 1C**. This was done for 11-31 days prior to all data reported here, and 27-56 days prior to perfusion and histological analyses). A 200 mW, 473 nm blue laser (VA-II-200-473, Optotronics) delivered stimulation controlled by a custom signal generator for 40 Hz or sham. The laser was calibrated using Thor Labs Optical Power Meter (PM100USB with S120C sensor) so that optogenetic stimulation was approximately 1.2 mW. Note that we calibrated the ferrule before implantation, but could not calibrate again after implantation but only estimate with a dummy ferrule, so there was likely some power loss for each mouse.

### Recording procedure

After recovery from surgery, daily recording and training sessions commenced. Each daily recording session included a 50-70 min sleep session, followed by 20 min of task (real or virtual), followed by another 50-70 min sleep session, and then a 1 hour stimulation session, either 40 Hz or sham (**Fig. 1D**. An electrode interface board (EIB-72-QC-Small, Neuralynx) was attached with a custom adapter to the recording array (Chang et al., 2013) with independently drivable tetrodes connected via a pair of unity-gain headstages (HS-36, Neuralynx) to the recording system (Digital Lynx SX, Neuralynx). Tetrodes were referenced to a tetrode wire in the corpus callosum and advanced as needed, up to 124 µm/day, while monitoring the audio and visual signal of the LFP and unit activity. Adjustments were made after a given day’s recording to allow stabilization overnight. Thresholded (adjusted prior to each session) spike waveforms were bandpass-filtered 0.6-6000 Hz and digitized at 32 kHz. A continuous trace was simultaneously collected for processing as LFP from one of the tetrode wires (bandpass-filtered 0.1-1000 Hz and digitized at 6400 Hz) and referenced to an electrode in corpus callosum. Mouse position was tracked using a colored dome of reflective tape for the real maze (or virtual position for the virtual maze), and live position information was used to trigger mfb stimulation rewards. Video-tracking or virtual position data was collected at 30 FPS and co-registered with spikes, LFPs, and event timestamps.

Spike data were automatically overclustered using KlustaKwik and manually adjusted using a modified version of MClust (A.D. Redish). All spike waveforms with a shape and across tetrode cluster signature (suggesting that they were likely MUA and not noise) were manually selected and merged into a single MUA cluster, as in (Harris et al., 2000; Wilber et al., 2017). Thus, MUA clusters included both well-isolated single units and poorly isolated single units (http://klustakwik.sourceforge.net; Harris et al., 2000; Cushing et al., 2020). LFP analyses were performed using custom-written MATLAB code (MathWorks, Natick, MA) or Freely Moving Animal (FMA) Toolbox (http://fmatoolbox.sourceforge.net/). The LFP signal was collected at 6400 Hz and subsequently resampled to 2000 Hz for further analysis, using the MATLAB *resample* function. Histology. At the conclusion of the experiment, marking lesions were made by applying a 5 µA current for 10 sec between each tetrode wire and the tail. One week later, mice were given an intraperitoneal injection of Euthasol and then transcardially perfused with 1X phosphate-buffered saline (PBS), followed by 4% paraformaldehyde (PFA) in PBS. The whole head was post-fixed for 24 hour to allow for easy identification of the tract representing location of tetrodes and mfb electrodes, and then the brain was removed and post-fixed for another 24 hours. Last, the brain was cryoprotected in a 30% sucrose solution in PBS. Frozen sections were cut coronally with a sliding microtome at a thickness of 40 µm and split into 6 evenly spaced series.

#### MOC78

Sections were rinsed twice in TBS then blocked in TBS containing 0.3% Triton X and 3% goat serum. Sections were incubated for one day with primary antibodies anti-mOC78 (M78, 0.3 μg/ml, monoclonal, rabbit, Abcam 205341) and anti-NeuN (1:1000, polyclonal, chicken, Millipore ABN91) in PBS containing 0.3% Triton-X. Next, sections were rinsed with TBS containing 0.3% Triton-X three times. Sections were then incubated with secondary antibodies anti-rabbit-alexa-488 (1:500, polyclonal, goat) and anti-chicken-alexa-594 (1:1000, polyclonal, goat) in TBS containing 0.3% Triton-X overnight. Finally, sections were rinsed once with TBS containing 0.3% Triton-X, and twice with TBS. Sections were mounted with a glycerol based mounting media containing DAPI and coverslipped then imaged.

#### MOC22

Procedure was the same as above, but with primary antibodies anti-mOC22 (M22, 0.5 μg/ml, monoclonal, rabbit, Abcam 205339) and anti-NeuN (1:1000, polyclonal, chicken, Millipore ABN91) in TBS containing 0.3% Triton-X. Secondaries, rinses, and mounting were the same as above.

#### Iba1

Procedure was the same as above, but with primary antibodies anti-Iba1 (1:1000, rabbit, Wako Chemicals, 019-19741) and anti-NeuN (1:1000, polyclonal, chicken, Millipore, ABN91) overnight, followed by secondary antibodies, anti-rabbit-alexa-488 (1:1000, polyclonal, goat) and anti-chicken-alexa-594 (1:1000, polyclonal, goat) respectively overnight. Sections were rinsed and mounted onto slides as described above.

#### NeuN

Procedure was the same as above, but with primary antibody anti-NeuN (1:1000, polyclonal, chicken, Millipore, ABN91) overnight, followed by secondary antibody anti-chicken-alexa-594 (1:500, polyclonal, goat) overnight. Sections were rinsed and mounted onto slides as described above. This was done to confirm virus expression in HPC.

#### Phosphorylated tau

Procedure was the same as above, but with primary antibodies anti-phospo-tau AT8 (1:500, monoclonal, mouse, Thermo Scientific MN1020) and anti-NeuN (1:1000, polyclonal, chicken, Millipore, ABN91) overnight followed by secondary antibodies, anti-mouse-alexa-488 (1:1000, polyclonal, goat) and anti-chicken-alexa-594 (1:500, polyclonal, goat) respectively, overnight. Sections were rinsed and mounted onto slides as described above.

#### GFAP and 6e10

Sections were rinsed twice in TBS then blocked in TBS containing 0.3% Triton X and 3% goat serum. Sections were incubated in anti-GFAP (1:2000, rabbit, Abcam, 7260), anti-6E10 (1:1000, β-amyloid 1-16, mouse, BioLegend, 803015), and anti-NeuN (1:1000, polyclonal, chicken, Millipore, ABN91) for two days was followed by secondary antibodies, anti-rabbit-alexa-594 (1:1000, polyclonal, goat), anti-mouse-alexa-488 (1:1000, polyclonal, goat), and anti-chicken-CF-350 (1:2000, polyclonal, goat) respectively overnight. Sections were rinsed and mounted onto slides, as described above.

### Image Acquisition

Whole slides were scanned using a scanning microscope at 10x magnification, high resolution images (Zeiss Axio Imager M2). Positive cells were automatically counted using Zeiss automatic segmentation tool (<u>ZEISS ZEN Intellesis</u>; M78, M22, 6e10, pTau, Iba1, and GFAP).

### Genotyping

We received homozygous 3xTg-AD mice from Dr. Frank LaFerla’s lab and have maintained our own breeding colony with occasional backcrosses with additional mice from the LaFerla lab. We confirmed that all mice used in the experiment contained each transgene using conventional PCR. DNA was extracted from the tails of each mouse. Homozygosity was confirmed by cutting the PS1 PCR fragment with the *BstEII* restriction enzyme. Only the mutated human PS1 gene contains a *BstEII* cut site and will be cut. The absence of an uncut PCR product indicated that the mouse was indeed homozygous for the human PS1. The presence of overexpressed APP and Tau were also confirmed by PCR. The previously published primers were used for amplifying the PS1 transgene, APP and Tau (Stimmell et al., 2019).

3xTg-AD mice were crossed with B6 PV^cre^ (Parvalbumin, 017320) from Jackson Labs. The F1 generation was crossed to create animals homozygous for both PV^cre^ and the 3 transgenes of 3xTg-AD (PS1, APP and Tau). This was accomplished by extracting DNA from the ear punch of each mouse and genotyping using conventional PCR. PCR primers and restriction digest for 3xTg-AD were used as described above. PV^cre^ primers used were provided by Jackson Labs (<u>Protocol 24678 - Pvalb<tm1(cre)Arbr>alternate2</u> (jax.org)). An additional backcross to 3xTg-AD was later performed.

## QUANTIFICATION AND STATISTICAL ANALYSIS

### Recording and Quantification

#### Sleep and LFP Analyses

First, still periods were extracted from the rest sessions as described previously (Euston et al., 2007; Wilber et al., 2017). The raw position data from each video frame was smoothed by convolution of both x and y position data with a normalized Gaussian function (standard deviation of 120 video frames). After smoothing, the instantaneous velocity was found by taking the difference in position between successive video frames. An epoch during which the velocity was constantly below 0.78 pixels/s (∼0.19 cm/s) for more than 2 min was considered a stillness period. All analyses of rest sessions were limited to these stillness periods. SWS and rapid eye movement (REM) sleep were distinguished using K-means clustering of the theta/delta power ratio extracted from the CA1 pyramidal layer LFP recorded during the stillness periods, as is common for assessing memory replay (**Fig. 1C**; Stan Leung, 1998; Buzsáki et al., 2003; Lansink et al., 2008; Sirota et al., 2008; Girardeau et al., 2009; Mizuseki et al., 2009; Mizuseki et al., 2011; Grosmark et al., 2012) and has been validated (Costa-Miserachs et al., 2003). Only SWS periods were included for the remaining sleep analyses, and only data sets with ≥10 min of sleep in both rest periods were included (**Table 1**). Delta wave troughs (DWTs), which correspond to cortical down states (Battaglia et al., 2004), were detected by digitally filtering the LFP trace from a representative cortical electrode in the 0.5-4 Hz range and detecting the peaks in the inverted signal that exceeded a mean +1.5 SD threshold. SWRs were detected from the HPC CA1 field LFP digitally filtered in the 75-300 Hz range, determined a priori, and consistent with a literature review of common frequency ranges (Maier et al., 2003; Nimmrich et al., 2005; Jadhav et al., 2016; Oliva et al., 2016; van de Ven et al., 2016; Roux et al., 2017; Yamamoto and Tonegawa, 2017). Events with peaks >6 SD above the mean and duration 20-100 ms were considered SWRs as in our previous work (Cushing et al., 2020). The SWR duration included the contiguous periods surrounding the peak and exceeding 2 SD above the mean. The SWR detection accuracy was visually validated on a subset of each analyzed dataset. This visual validation process caused us to adjust the SWR detection criteria for two mice (one 40 Hz, one sham stimulation). For these mice, events with peaks >7 SD were used instead. Making this adjustment ensured that a similar signal to noise ratio was applied to all mice. In order to assess HPC-PC interactions, SWRs were cross-correlated with DWTs, and the SWR count binned (bin size 16.67 ms - time bin of 100 ms divided by 6x compression factor) as a function of position in time from the DWT, then z-scored. Since timing of the peak SWRxDWT correlation sometimes appears in slightly different time bins across animals and sessions, the peak value was compared irrespective of the precise position in time. Sleep spindles were detected on the same cortical electrode as DWT’s. The raw LFP signal was digitally filtered in 8-18 Hz range, followed by rectifying and z-scoring of the resulting signal. Sleep spindles were defined as the time windows with peaks >3.5 SD above the mean and duration of 0.5-3 sec. Spindle duration was defined as the contiguous window around the spindle peak with envelope amplitude >2 SD above the mean. Spindle onset and offset were defined as the first and the last timepoints of individual spindle events. Raw LFP in the window +/-2 sec surrounding the detected DWT was bandpass-filtered in spindle range (8-18 Hz) and the peak of the filtered signal was located. The delta-spindle coupling phase was defined as the phase of delta wave that coincided with spindle peak. The circular mean of delta-spindle coupling phases was calculated for each individual sleep epoch, using the circ_mean.m function from MATLAB Circular Statistic Toolbox (Berens, 2009).

**Table 1.** Exclusion Criteria. Numbers of mice from each treatment group that have been eliminated for each analysis, as well as the reason for elimination.

| Analysis | Behavior | Sleep – still time | Sleep architecture and events | Pathology (HPC) | Pathology (PC) |
| --- | --- | --- | --- | --- | --- |
| # mice (40Hz, Sham) | 5,6 | 7,6 | 7,5 | 6,6 | 5-6, 5-6 |
| Reasons for exclusion | 2 40Hz mice excluded from RMANOVA due to missing velocity data for some days of behavior (included in figure and correlations where appropriate) | All mice included | 1 Sham mouse excluded due broken drive | 1 40Hz mouse excluded due to no stains of HPC | 1-2 40Hz mice and 1 Sham mouse excluded from some stains due to no PC present on those sections |

### Pathology Quantification

Images were loaded into Zeiss, and experimenters segmented the regions of interest (ROIs) using Zeiss segmentation tool. ROIs were defined using the Allen Brain Atlas (Oh et al., 2014). After ROIs were determined, thresholds for fluorescence for individual sections and ROIs were set. Fluorescence for NeuN was set first, followed by fluorescence for pathology. For glial cells, fluorescence was only thresholded for the cells. Finally, a threshold for size and roundness was applied. Neurons had to have greater area than 40 µm^2^, and had to have a circularity greater than 0.45. Glial cells had to have area greater than 10-20 µm^2^, with threshold set to include all cell bodies and processes but not any background for each image, and no threshold for circularity (Norden et al., 2016). Then, neurons were segmented using the NeuN color channel and counts of cells positive for each marker (M78, M22, 6e10, pTau, Iba1, and GFAP) were obtained automatically within each region of interest using Zeiss automatic segmentation tool (<u>ZEISS ZENIntellesis)</u>. Counts were performed on every other section from the 1:6 series for Aβ and tau stain (i.e., every 12^th^ section), and performed on each section for glial stains, which were stained on a 1:12 series. Cell number (glia) or cells positive for pathology were divided by the area to obtain proportions of positive cells.

### Statistical Analyses

Statistical comparisons were performed using two-way repeated measures ANOVAs (group x sleep session or genotype x day of behavior), paired t-tests (used when RMANOVA was not possible, for example due to imbalanced data sets, averaged within animal), and chi-square analyses. Correlations were run using R package correlation with multiple levels correcting for group membership. For all statistical analyses, p<0.05 was considered significant after a Bonferroni correction for multiple comparisons was applied (when appropriate). Bonferroni corrected critical values are only shown for Bonferroni corrections that shift an uncorrected significant p-value to non-significant and not when the statistical result remains significant after Bonferroni correction. Statistical analyses were performed using MATLAB Statistics and Circular Statistics toolboxes (Berens, 2009), StatView (SAS Institute Inc.), SigmaPlot (Systat Software Inc.), and R (correlation and cocor packages). Distributions of session-level circular means between groups for delta-wave phase relationship analyses were compared using the Watson-Williams test (ww_test function from MATLAB Circular Statistic Toolbox).

## Results

### Ameliorated spatial learning and memory deficits

The 40 Hz stimulation of PV+ interneurons in HPC resulted in an increase in 40 Hz power spectral density for 40 Hz mice (**Fig. 1F**), but not sham. Previously, 3xTg-AD mice were impaired on this task segment, showing less slowing in the reward zone that corresponded to deficits in HPC-PC coordination. We hypothesized that 40 Hz stimulation of HPC interneurons would result in the rescuing of this spatial navigation deficit. We assessed spatial navigation performance on the virtual maze during the first 6-days of the task when memory related brain dynamics are strongest as in (Cushing et al., 2020). We compared the z-scored velocity in the reward zone between the 40 Hz stimulated and sham stimulated 3xTg-AD/ PV^cre^ mice during the same time period. The 40 Hz mice slowed significantly more in the reward zone than sham stimulated mice (**Fig. 1G**, RMANOVA; F_(1,9)_=8.957, p=0.015, no day effect or interaction, Fs_(5,45)_≤0.784, ps≥0.566). To rule out the possibility that differences in motor abilities led to this impairment, we also compared velocity on the track at a significant distance from the reward zone to ensure motor differences did not explain the group difference, and found no difference between groups (F_(1,9)_=0.267, p=0.267, no day effect or interaction, Fs_(5,45)_≤1.378, ps≥0.2507). Thus, the 3xTg-AD/PV^cre^ mice that received 40 Hz stimulation to HPC demonstrated improved spatial navigation on the VM compared to sham stimulated mice.

### HPC-PC interactions are rescued by 40 Hz stimulation, but do not predict subsequent VM performance

We hypothesized that hippocampal-PC interactions during sleep would be rescued by 40 Hz stimulation, ameliorating the deficit we previously identified that could reflect impaired cross-structure coordination of memory replay (Cushing et al., 2020). As previously reported, we assessed PC-HPC coordination using SWRs in the HPC and DWTs in PC, and cross-correlating the two events in pre-task sleep and post-task sleep (**Fig. 2A**, strong NonTg example). 3xTg-AD mice previously demonstrated a lack of learning induced increase in SWR-DWT coordination, while NonTg control mice showed a significant increase in SWR-DWT correlations in post-task sleep compared to pre-task sleep (**Fig. 2B**; Cushing et al., 2020). We assessed the proportion of data sets with a higher SWR-DWT correlation in post-task sleep compared to pre-task sleep. The 40 Hz stimulated mice showed an increase in significantly more sessions (79%, **Fig. 2C**) compared to SHAM mice (47%, χ^2^ =4.39, p=0.036). We also assessed how much the average SWR-DWT peak value changed across sleep session using animal means between groups. NonTg mice previously had a significant increase in SWR-DWT peak correlations in post-task sleep (**Fig. 2D**, paired t-test; t_(4)_=3.69, p=0.02) while 3xTg-AD mice had a decrease in SWR-DWT correlations across sleep session (**Fig. 2E**, paired t-test; t_(6)_=-2.8, p=0.04). 40 Hz stimulation reversed this impairment as 40 Hz mice had a significant increase in SWR-DWT peak correlations (**Fig. 2G**, paired t-test; t_(6)_=5.26, p=0.002), while sham mice did not (**Fig. 2H**, paired t-test; t_(4)_=0.89, p=0.424). In order to assess if there was a difference in the distributions of SWRxDWT during the 1 sec surrounding the DWT between groups, we performed a KS test on post – pre distributions between stimulation groups. We found there was a significant difference between the two distributions (D=0.2333, p=0.0381).

**Figure 2.**
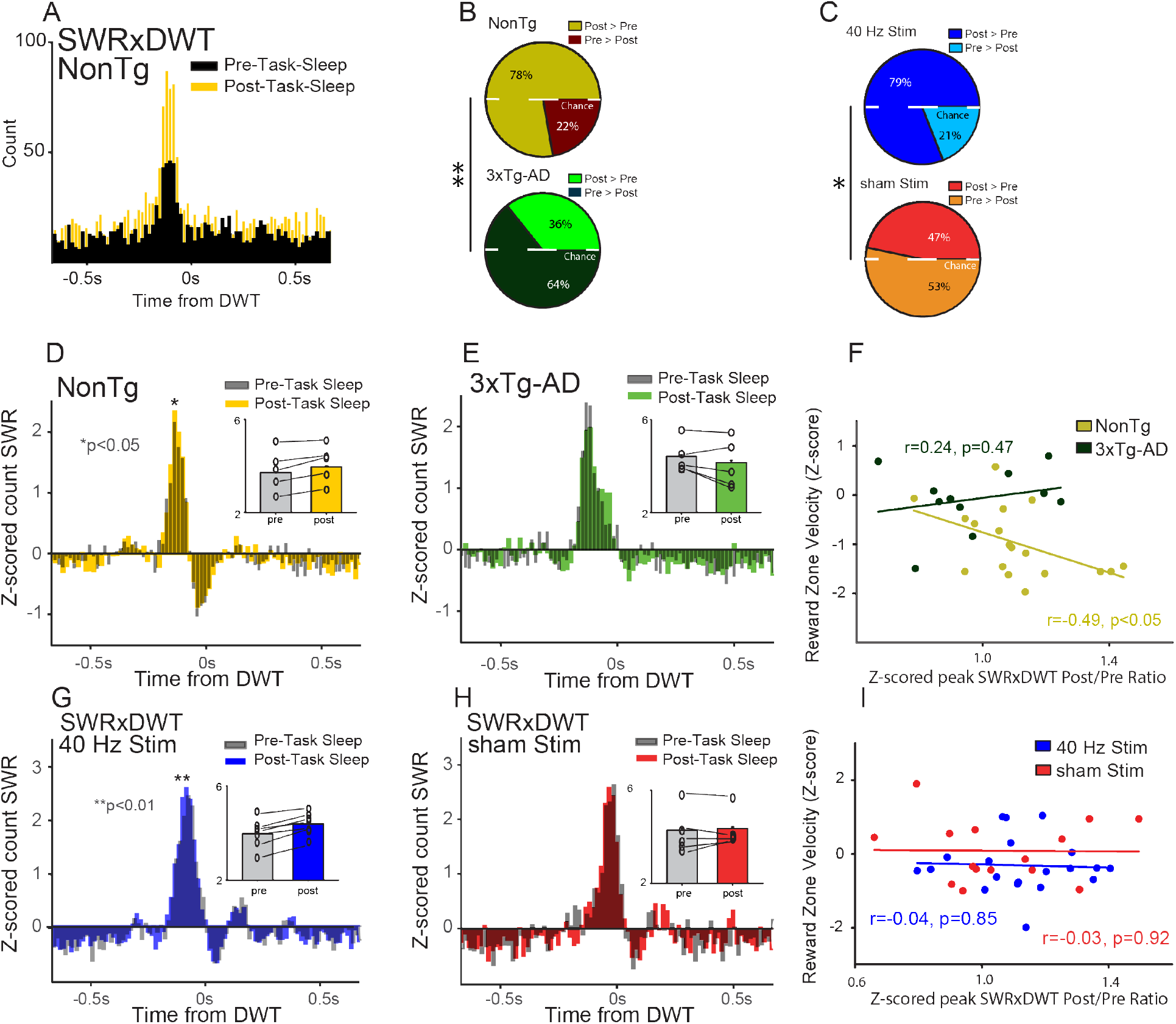
PC-HPC network coordination is rescued by 40 Hz HPC stimulation, but is dissociated from behavior. **A.** Strong example cross-correlation of SWR and DWTs from NonTg mice. SWRs tend to occur immediately preceding the DWT and this relationship is strengthened in post-task sleep (yellow) relative to pre-task sleep (black). **B**. The proportion of data sets in which the cross-correlation peak was larger in post-task sleep was significantly higher in NonTg mice (gold/garnet) than 3xTg-AD mice (light/dark green). The HPC-PC reactivation event coupling was significantly reduced in 3xTg-AD mice compared to NonTg mice (χ2(1)=5.77, p≤0.01). **C**. The proportion of data sets in which the cross-correlation peak was larger in post-task sleep was significantly higher for 40 Hz stim mice (light/dark blue) than sham mice sets (orange/red). The HPC-PC reactivation event coupling is reduced in sham mice for post-task relative to pre-task sleep (χ2(1)=3.88, p<0.05). **D**. Z-scored count of SWR occurring relative to the DWT for NonTg mice. The temporal relationship between SWRs and DWTs was significantly strengthened in post-task sleep (yellow) relative to pre-task sleep (grey). *Inset* is the average peak SWRxDWT for pre and post-task sleep. Note, that this number is higher than shown on the main graph as the peak moves bins between data sets, resulting in the total average being lower. Data from the same mouse is connected with a line. **E**. Same as D for 3xTg-AD mice. SWRs tended to precede the DWT, but this relationship was not strengthened in post-task sleep (green) relative to pre-task sleep (grey). **F**. The relationship between SWR-DWT coupling and performance on the VM the subsequent day. For NonTg mice, the greater the strengthening of SWR-DWT coupling in post-task sleep relative to pre-task sleep, the more the mice slowed down in the reward zone the following day (yellow, r=-0.49, p<0.05). 3xTg-AD mice showed no such relationship (green, r=0.24, p<0.47). **G**. Mean Z-scored cross-correlation between HPC SWR and PC DWT for 40 Hz stimulated mice. SWRs precede DWT by about 117 ms, and this relationship is strengthened in post-task (blue, n=7) versus pre-task -sleep (grey) in 40 Hz stimulation. **H**. Same as G for sham stimulated mice, which also have a peak with SWRs again preceding DWTs by about 83 ms, but the peak does not increase in post-task sleep (red, n=5) compared to pre-task sleep (grey). **I**. in both 40 Hz and sham stimulated mice, DWT-SWR coupling is not predictive of improved behavioral performance the subsequent day. *p<0.05, ** p<0.01. Panels A, B, D-F adapted from (Cushing et al., 2020).

In NonTg mice studied previously, but not 3xTg-AD mice, the increase in the peak temporal correlation of SWR-DWT in post-task sleep over pre-task sleep predicted improved performance on the virtual maze the following day (**Fig. 2F**; Cushing et al., 2020). However, despite the restored PC-HPC coupling, the coupling strength did not predict performance on the virtual maze the following day for 40 Hz mice (**Fig. 2I**, r_(18)_=-0.04, p=0.851) or sham mice (r_(14)_=-0.03, p=0.927). Thus, despite rescuing both the SWR-DWT temporal relationship and performance on the virtual maze, 40 Hz stimulation did not restore the relationship between mesoscale brain activity patterns and behavior, suggesting that prior pathology accumulation may have decoupled this cortico-HPC coupling from behavior.

### Unchanged PC interactions

It has been suggested that the temporal relationships between SWRs, DWTs, and spindles contributes to memory consolidation (Clemens et al., 2007; Jiang et al., 2019). Previously, we found that there was a strong temporal relationship between SWRs and spindles in both NonTg and 3xTg-AD mice (Cushing et al., 2020). We once again found a strong temporal relationship between SWRs, spindles, and DWTs, with SWRs tending to follow the spindle, and DWTs following both spindles and SWRs. We performed the same cross-correlation analysis as in SWR-DWT for SWR-spindle correlations, and found there was no difference in the increase in temporal relationship for either 40 Hz stimulated mice (**Fig. 3** *Top*, paired t-test; t_(6)_=0.177, p=0.8655), or sham stimulated mice (t_(4)_=0.176, p=0.8691).

**Figure 3.**
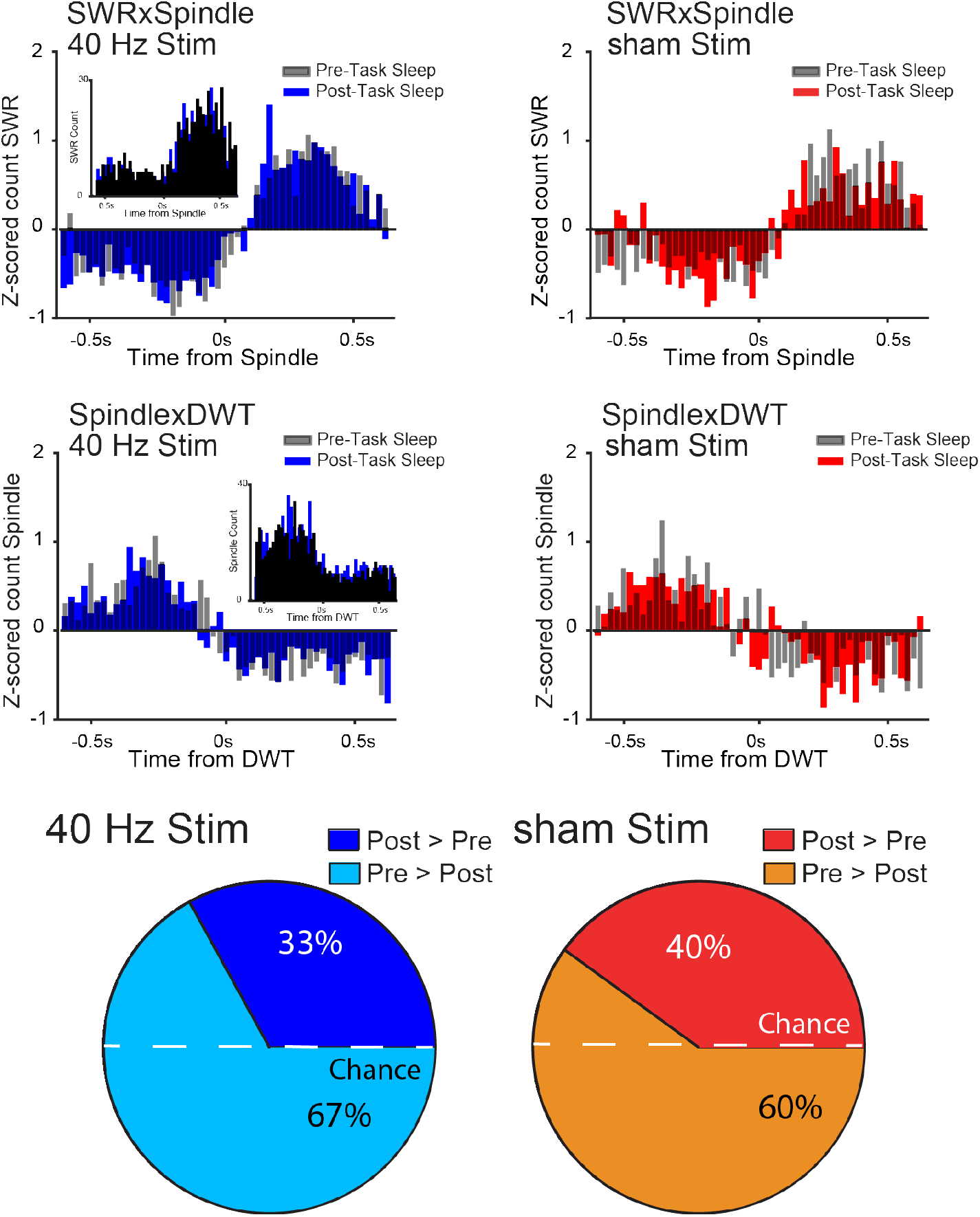
Spindle-DWT coupling is unaffected by 40 Hz HPC stimulation. *Top*. Mean Z-scored cross-correlation between HPC SWR and PC spindles. We found that for 40 Hz stimulated mice (*left*) SWRs follow spindles in both pre-task sleep (grey) and post-task sleep (blue, n=7). Similarly, for sham stimulated mice (*right*) SWRs followed spindles in post-task sleep (red, n=6) and pre-task sleep (grey). *Middle.* Mean Z-scored cross-correlation between PC spindles and DWT for 40 Hz stim mice (*left*) and sham stimulated mice (*right*). Spindles tend to precede the DWT in both pre-task-sleep (grey) and post-task sleep (blue, n=7) in 40 Hz mice and sham stimulated mice (*right*). *Bottom*. The proportion of data sets in which the cross-correlation peak DW-spindle was larger in post-task sleep was not significantly different for 40 Hz stim mouse data sets (light/dark blue) than sham mouse data sets (orange/red, χ2(1)= 0.18, p=0.673).

We also had previously identified deficits in spindle-DWT relationship in 3xTg-AD mice (Cushing et al., 2020). We assessed spindle-DWT correlations in a similar manner to spindle-SWR. We did not expect to see an effect of stimulation on the correlation of spindles and DWT, as neither event originates in the hippocampus. Spindles tended to precede the delta waves (**Fig. 3** *Middle*). We assessed if there was a change in spindle-DWT correlations by comparing the peak of this relationship in pre-task and post-task sleep. We found that 33% of data sets that showed an increase in post-task sleep for 40 Hz stimulated animals (**Fig. 3** *Bottom*). This was not significantly different from sham stimulated animals, which showed an increase in 40% of data sets (χ^2^ =0.18, p=0.673). 40 Hz stimulated mice did not show a significant difference from pre-task to post-task sleep (paired t-test; t_(6)_=1.22, p=0.2683). Sham mice also did show a difference between pre-task and post-task sleep (paired t-test; t_(4)_=1.648, p=0.1747). We also identified deficits in the spindle-DWT relationship in 3xTg-AD mice, with spindle power no longer being strongest during the DWT peak, but being distributed across the DW (Cushing et al., 2020). When we assessed spindle-DW relationships in 40 Hz and sham stimulated mice, we found that spindles tended to occur at the peak of the DW, similar to NonTg mice, and that there was no significant difference between stimulation (F_(1,78)_=0.747, p=0.3902). This indicates that HPC 40 Hz stimulation does not have a significant impact on spindle-Delta or spindle-SWR coupling different from sham stimulation. Interestingly, we previously found that in NonTg mice the spindle-DWT correlation is larger in post-task sleep in 70% of the data sets, suggesting that 40 Hz stimulation targeting hippocampus may not fully restore brain thalamocortical coordination (Cushing et al., 2020). Such a finding is not unexpected since 40 Hz stimulation was not delivered to thalamus.

### Individual Markers of Memory Related Events Predict Subsequent Behavioral Performance in 40 Hz Stimulated Mice

SWRs in HPC, as well as DWTs and thalamocortical spindles in PC, are important events for memory (Clemens et al., 2006; Eschenko et al., 2006; Diba and Buzsáki, 2007; Diekelmann and Born, 2010; Roux et al., 2017; Fernández-Ruiz et al., 2019b; Kim et al., 2019; Sanchez-Aguilera and Quintanilla, 2021). We assessed both the number and rate (number/second of SWS) of each event between 40 Hz and sham stimulation groups. SWR number and rate did not differ across stimulation type (**Fig. 4** *Top*, Fs_(1,10)_ ≤0.457, ps≥0.5145, no sleep session effect or interaction, Fs_(1,10)_≤3.521, 0.0901). Likewise, DWT number and rate did not differ (Fs_(1,10)_ ≤0.936, ps≥0.3561, no sleep session effect or interaction, Fs_(1,10)_≤3.777, 0.0806), nor did spindle number (F_(1,10)_ =0.311, p=0.5892, no sleep session effect or interaction, Fs_(1,10)_≤0.886, 0.3688). Spindle rate also did not differ between groups (F_(1,10)_=2.583, p=0.1391), but did increase in post-task sleep (not shown; F_(1,10)_ =5.296, p=0.0442, no interaction F_(1,10)_=0.22, p=0.6493), which is expected given that post-task sleep is thought to be critical for memory formation.

**Figure 4.**
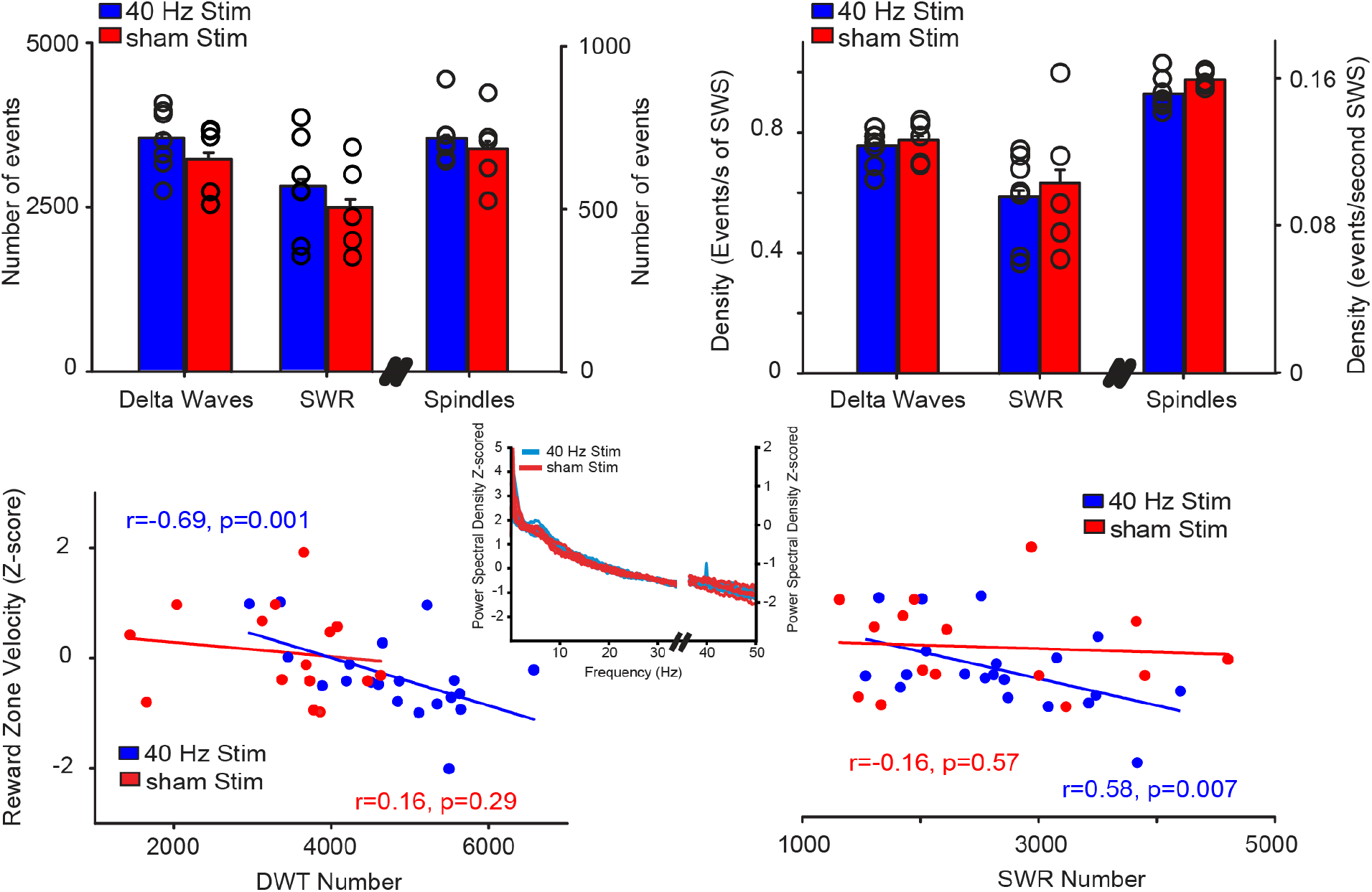
DWT and SWR number predict subsequent performance on the VM in 40 Hz but not sham stimulated mice. *Top Left*. The mean (±SEM) number of memory reactivation markers; PC DWT, HPC SWR, and thalamocortical spindles. 40 Hz stimulated mice (blue) showed no difference in density of markers of memory related brain dynamics compared to sham stimulated mice (red). *Top Right*. Density (mean ±SEM) of memory related activity markers. No differences between 40 Hz stim mice or sham stim mice in density of events. *Bottom Left.* The total number of DWT predicted behavioral performance in 40 Hz stim mice, but not sham mice. *Bottom right*. The total number of SWR predicted behavioral performance in 40 Hz stim mice, but not sham mice. Inset shows example PC power spectral density during stimulation (40 Hz or sham) for each mouse. Most 40 Hz stimulated mice had increases in 40 Hz power in PC during stimulation session.

Given that the temporal correlation between pairs of SWR, DWT, and Spindle events (e.g., SWR-DWT) either did not vary across stimulation type (Spindle-SWR, Spindle-DWT) or did not predict subsequent performance (SWR-DWT) for either 40 Hz or sham stimulated mice, we hypothesized that markers of memory related events in individual brain regions could predict performance instead. We assessed whether the number or rate of these markers of memory related activity predicted performance on VM the subsequent day for either 40 Hz or sham stimulated mice. Increased number of SWRs predicted improved performance on the maze the following day for 40 Hz stimulated mice (r_(18)_=-0.58, p=0.007), but not sham stimulated mice (r_(14)_=-0.16, p=0.565) (Fisher r-to-z, z=-2.1573, p=0.0155). SWR rate did not predict subsequent performance for either 40 Hz stimulated mice (r_(18)_=-0.07, p=0.763) or sham stimulated mice (r_(14)_=0.13, p=0.636), suggesting that the number of events may be the critical driver of performance. Similarly, the number of DWT events predicted performance on the VM the following day for 40 Hz stimulated mice (r_(18)_=-0.69, p=0.001), but not sham stimulated mice (r_(14)_=-0.16, p=0.552) (z=-1.7979, p=0.0361). DWT rate did not correlate with subsequent performance for either 40 Hz stimulated (r_(18)_=-0.04, p=0.876) or sham stimulated mice (r_(14)_=0.02, p=0.927). Neither spindle number or rate predicted subsequent performance on the VM for 40 Hz stimulated mice (rs_(18)_ ≤│0.34│, ps≥0.143) or sham stimulated mice (rs_(14)_ ≤│0.15│, ps≥0.591). Our findings that spindle rate and number do not predict performance combined with our findings that SWR-DWT coupling also do not predict subsequent performance are consistent with other studies suggesting that spindles mediate HPC-cortical coupling during SWRs (Ngo et al., 2020).

#### Amount of slow wave sleep predicts performance on the virtual maze in 40 Hz mice only

We hypothesized that 40 Hz stimulation might result in changes in sleep dynamics, as 40 Hz multisensory stimulation has been shown to reduce sleep disturbances in AD patients (Cimenser et al., 2020). AD has been reported to reduce and fragment circadian sleep, and we previously identified changes in sleep during the rest sessions employed in our study design in 6-month female 3xTg-AD mice (e.g., increased time still proportion of still-time spent in SWS; Roh et al., 2012; Cushing et al., 2020). Further, multisensory 40 Hz stimulation has been reported to increase sleep compared to sham stimulation (Cimenser et al., 2020). Thus, we first assessed basic sleep characteristics in 40 Hz optogenetically stimulated mice. We found that 40 Hz mice and sham mice did not differ in still-time (**Fig. 5**, F_(1,11)_=0.157, p=0.699, no day effect or interaction, Fs_(1,11)_≤1.01, ps≥0.337). In order to control for any variations in sleep session length, we also assessed stillness as a proportion of sleep session length, and found 40 Hz mice and sham mice again did not differ (F_(1,11)_=0.416, p=0.532, no sleep session effect or interaction, Fs_(1,11)_≤0.766, ps≥0.400). Next, because our measures of brain dynamics are limited to data sets with sufficient sleep (≥10 min) and it is not necessary to limit to these data sets for these basic sleep measures, we also analyzed basic sleep characteristics for all data sets (including those excluded for previous analyses). There were again no differences in time or proportion of time spent still (Fs_(1,11))_≤0.654, ps≥0.4357, no sleep session effects or interactions, Fs_(1,11)_≤1.106, ps≥0.3352).

**Figure 5.**
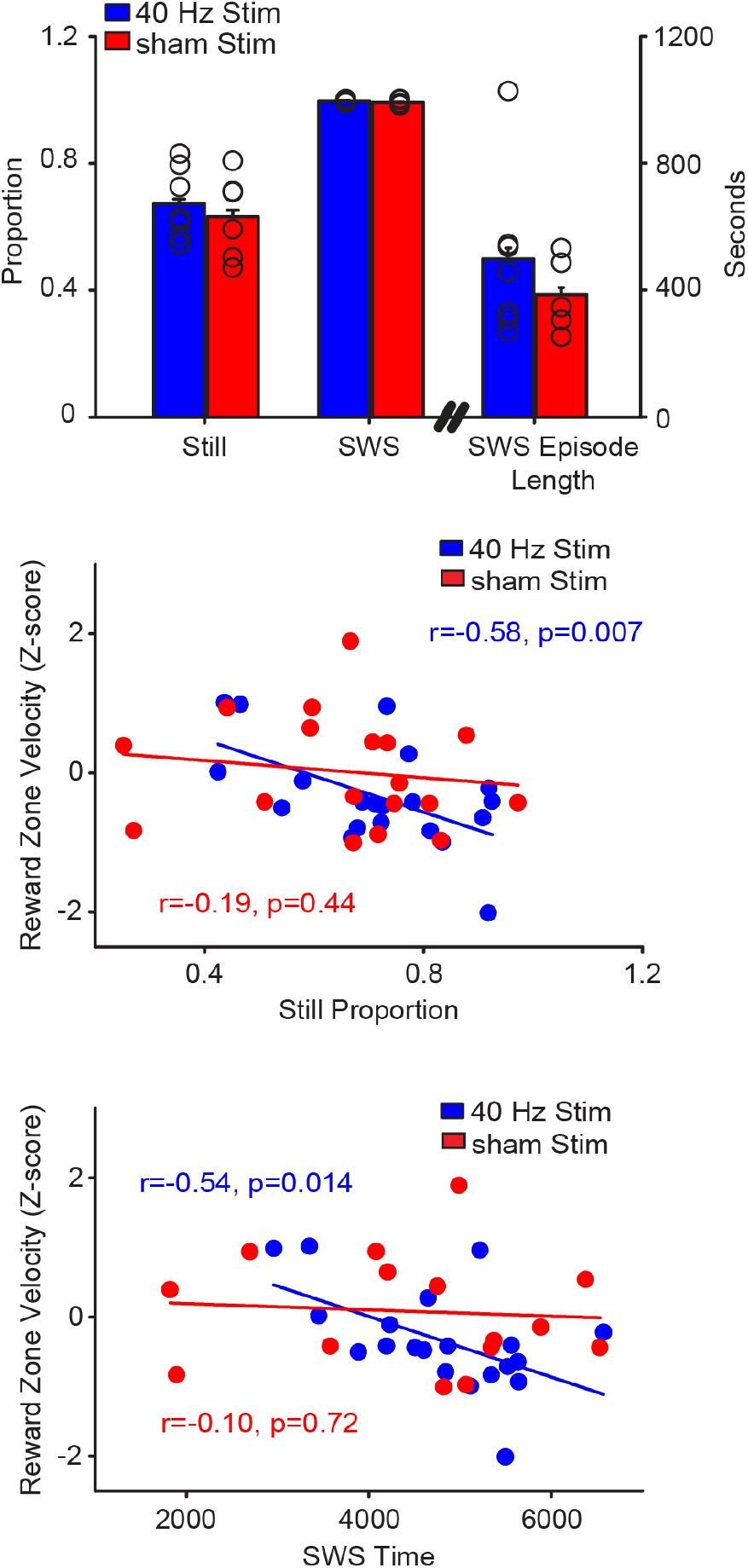
Still-time and SWS predict subsequent performance in 40 Hz mice. *Top*. Mean (±SEM) proportion of stillness, slow wave sleep (SWS), and SWS episode length for 40 Hz (blue), and Sham stimulated mice (red). Stimulation did not result in any changes in time spent in sleep or SWS. *Middle*. Proportion of sleep sessions spent still predicted slowing in the reward zone the following day in 40 Hz (blue), but not Sham stimulated mice (red). *Bottom*. Time spent in SWS predicted slowing in the reward zone the following day in 40 Hz (blue), but not Sham stimulated mice (red).

We also assessed time and proportion of time spent in REM and SWS and found that 40 Hz mice and sham mice spent equivalent time in SWS (F_(1,10)_=1.746, p=0.2159, no sleep session effect or interaction, Fs_(1,10)_≤0.745, ps≥0.4084). The proportion of total sleep time also did not differ between groups (F_(1,10)_=0.940, p=0.3551, no sleep session effect or interaction, Fs_(1,10)_≤1.923, ps≥0.1956). Number of episodes of SWS and length of SWS episodes also did not differ between groups (Fs_(1,10))_≤0.763, ps≥0.0.403, no sleep session effects or interactions, Fs_(1,10)_≤0.631, ps≥0.4455). Likewise, there was no difference in time spent in REM between stimulation groups (F_(1,10)_=0.281, p=0.6074, no sleep session effect or interaction, Fs_(1,10)_≤3.633, ps≥0.0858), or in REM as a proportion of total sleep time (F_(1,10)_=0.905, p=0.3639, no sleep session effect or interaction, Fs_(1,10)_≤1.360, ps≥0.2706).

After comparing how much 40 Hz and sham mice slept, we looked at how the amount of sleep correlated with performance on the virtual maze the following day. We hypothesized that increased time spent in SWS could predict performance on the maze the subsequent day, as numbers of both SWRs and DWTs predicted subsequent performance. We found that the more time 40 Hz stimulated mice spent still during sleep sessions, the more they slowed down in the reward zone the following day (r_(18)_=-0.54, p=0.013). Still-time did not predict performance for sham stimulated mice (r_(14)_=-0.18, p=0.466, Fisher r-to-z, z=-1.1055, p=0.1345). Furthermore, amount of time spent in SWS specifically predicted performance on the virtual maze the following day for 40 Hz stimulated mice (r_(18)_=-0.54, p=0.014), but not sham stimulated mice (r_(14)_=-0.10, p=0.723; Fisher r-to-z, z=-1.3193, p=0.0935). SWS episode number and SWS episode length did not correlate with performance the following day for 40 Hz mice (rs_(18)_ ≤│0.30│, ps≥0.201) or sham mice (rs_(14)_ ≤│0.19│, ps≥0.47). Thus, time spent in SWS predicted behavioral performance the following day in 40 Hz stimulated mice only.

Since the number of both SWRs and DWTs predicted performance on the the VM the following day, we wanted to make sure this was not just a statistical artifact of SWS time predicting performance and also driving higher numbers of these events. Thus, we ran partial correlations between performance and SWRs and DWT numbers, controlling for the effect of SWS time. For 40 Hz mice, SWR number no longer predicted subsequent performance (r_(18)_=-0.40, p=0.08), but DWT number did (r_(18)_=-0.51, p=0.02). For sham mice, neither SWR nor DWT number predicted subsequent performance (rs_(14)_ ≤│0.19│, ps≥0.48; Fisher r-to-z (DWT only) r=-0.9699, p=0.1660). Thus, DWT number predicts subsequent behavioral performance even when controlling for sleep.

#### Tau and Aβ relationships with Brain Dynamics and Behavior

40 Hz optogenetic, multisensory, and light only stimulation has previously been shown to result in a reduction in Aβ (Iaccarino et al., 2016; Singer et al., 2018; Martorell et al., 2019), though some studies have not confirmed this finding when using light only stimulation (Soula et al., 2023; Yang and Lai, 2023). We hypothesized that it would be difficult to see substantial clearance of Aβ or tau. This is because in the lightweight recording array we use, tetrodes become stuck by the end of the experiment and cannot be retracted from the brain before the array is removed, meaning that brain material around the recording array is lost (see **Fig. 6K** for examples of drive extraction damage). Similarly, the more intact hemisphere where stimulation was applied has fluorescently labeled AAV again making the automated pathology quantification we employed here essentially impossible (**Fig. 6A**). The surgical implantation of the recording array would also result in an immune response that we expected would make it difficult to see impacts of 40 Hz stimulation on microglia. Nonetheless, we assessed the impact of 40 Hz stimulation to HPC on Aβ and tau in the regions with recording data (HPC & PC) for the remaining intact tissue on the hemisphere with the recording array (contralateral to stimulation). We quantified the density of neurons with intracellular phosphorylated tau (pTau), Aβ 1-16 (6e10), and Aβ 1-42 (M22 and M78) and compared between groups (**Fig. 6**). There was no difference between 40 Hz stimulated mice and sham stimulated mice in pTau+ cell density in PC (**Fig. 6B-C**, t_(8)_ =0.375, p=0.7176), or dCA1 and vCA1 (ts_(10)_≤1.443, ps≥0.1795). However, 40 Hz stimulated mice also showed a significant reduction in M22+ cell density in PC compared to sham stimulated mice (**Fig. 6D-E**, t_(8)_ =3.536, p=0.0077), but not in dCA1 or vCA1 (ts_(10)_≤1.035, ps≥0.3258). Similarly, 40 Hz stimulated mice had a significant reduction in M78+ cell density in PC compared to sham stimulated mice (**Fig. 6F-G**, t_(9))_=2.182, p=0.05), but not in dCA1 or vCA1 (ts_(9)_≤1.536, ps≥0.1589). Finally, 40 Hz stimulated mice had a significant reduction in 6e10+ cell density in vCA1 relative to sham stimulated mice (**Fig. 6B-C** *Left*, unpaired t-test, t_(10)_ =2.884, p=0.0163), but not in dCA1 or PC (ts_(10)_≤1.735, ps≥0.1135).

**Figure 6.**
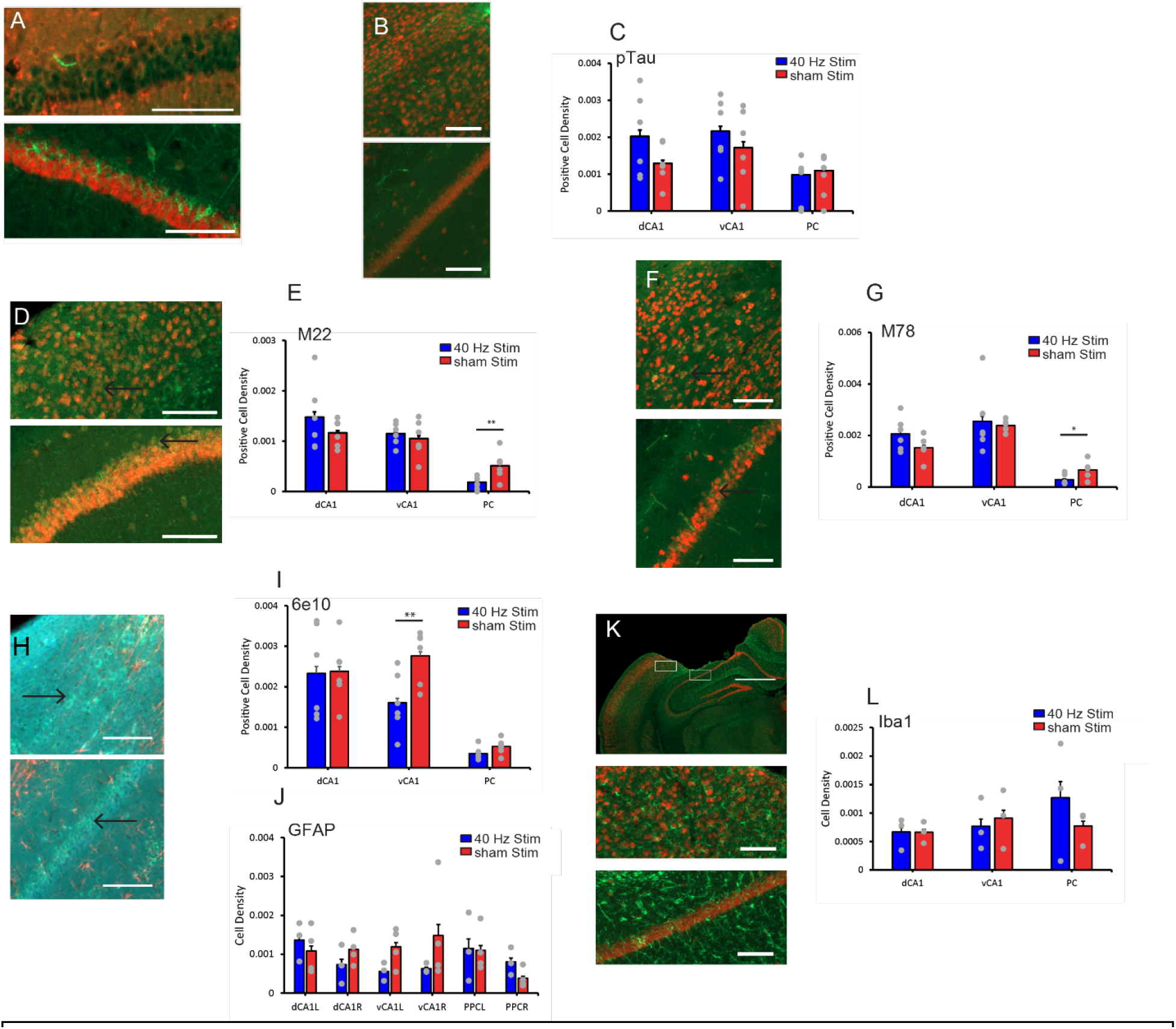
40 Hz HPC stimulation has modest impacts on pathology but not glia in the PC-HPC network. **A.** *Top*. Example image of fluorescent parvalbumin (red) showing which cells in HPC are parvalbumin positive. *Bottom*. Example image of NeuN (red) staining showing cells expressing AAV (green) in HPC PV cells. AAV expression is apparent along the dorsal border of the dorsal CA1 field of the HPC, the same location where parvalbumin positive cells are observed. **B**. Example staining of pTau (green) and NeuN (red), with PC (top) and HPC (bottom). **C.** Mean ± SEM positive cell density for pTau for 40 Hz (blue) and sham (red) stimulated mice for right hemisphere. **D.** Same as B for M22 (green) and NeuN (red). **E.** Same as C for M22, which showed a reduction in positive cell density in PC with a reduction in 40 Hz mice. **F.** Same as B, for M78 (green) and NeuN (red). **G.** Same as C for M78, which showed a reduction in positive cell density in PC of 40 Hz mice. **H.** Example staining of GFAP (red), 6e10 (green) and NeuN (blue) for PC (top) and HPC (bottom). Note, because GFAP can be labeled with a red fluorophore it can be quantified in both hemispheres. **I**. Mean ± SEM positive cell density for 6e10 for 40 Hz (blue) and sham (red) stimulated mice for right hemisphere. 40 Hz stim resulted in a decrease in 6e10+ cell density in vCA1 only. **J**. Same as C for astrocytes using GFAP, for both hemispheres. **K**. Same as I but for microglia using Iba1 (green) and NeuN (red). **L**. Same as C for Iba1. All scale bars on hemispheres are 1000μm, and all scale bars on insets are 100μm. Arrows show examples of positive cells.

Next, we correlated Aβ and tau load in the regions with recording data (HPC and PC) with performance on the virtual maze. 6e10+ cell density did not correlate performance on the maze for PC, dCA1, or vCA1 (multilevel correlation with stimulation as factor, **Table 2**, rs_(10)_ ≤│0.47│, ps≥0.119). Similarly, M22+ cell density did not correlate with performance on the maze for PC (r_(9)_ =0.50, p=0.115), dCA1 or vCA1 (rs_(9)_ ≤│0.54│, ps≥0.067).

**Table 2.**
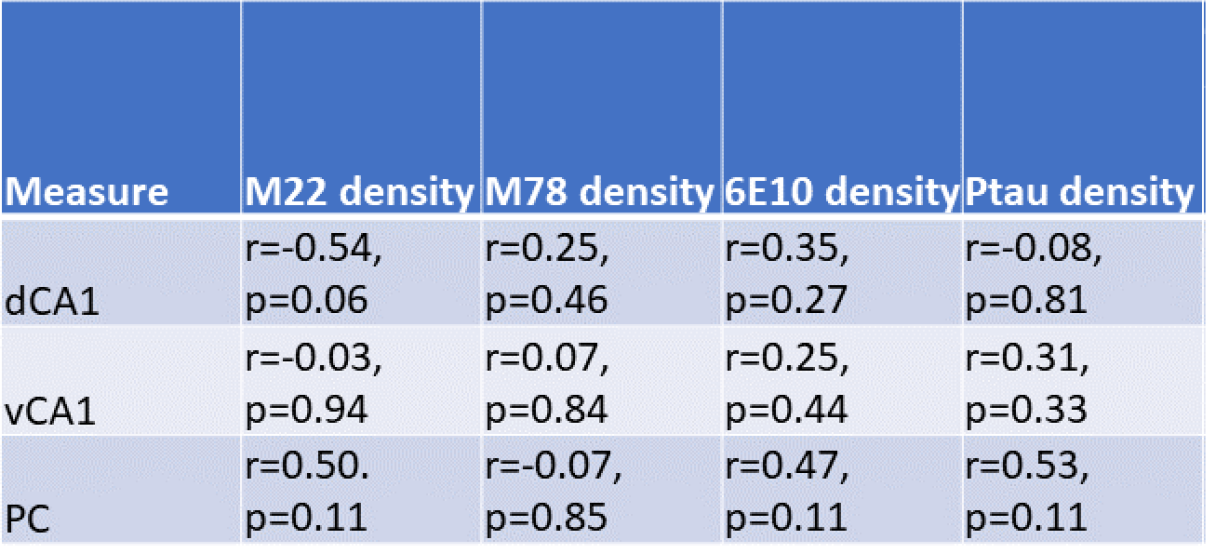
Relationship between positive cell density and measures of behavioral performance. R and p values for correlations between M22+ cells, M78+ cells, 6e10+ cells, and pTau+ cells in dorsal CA1 (dCA1), ventral CA1 (vCA1), and PC and slowing in the reward zone.

| Measure | M22 density | M78 density | 6E10 density | Ptau density |
| --- | --- | --- | --- | --- |
| dCA1 | r=-0.54,<br>p=0.06 | r=0.25,<br>p=0.46 | r=0.35,<br>p=0.27 | r=-0.08,<br>p=0.81 |
| vCA1 | r=-0.03,<br>p=0.94 | r=0.07,<br>p=0.84 | r=0.25,<br>p=0.44 | r=0.31,<br>p=0.33 |
| PC | r=0.50,<br>p=0.11 | r=-0.07,<br>p=0.85 | r=0.47,<br>p=0.11 | r=0.53,<br>p=0.11 |

Similarly, M78+ cell density also did not correlate with performance on the maze for PC, dCA1, or vCA1 (rs_(9)_ ≤│0.25│, ps≥0.458). Finally, pTau+ cell density did not correlate with performance on the maze for PC (r_(8)_ =0.53, p=0.116), or dCA1 or vCA1 (rs_(10)_ ≤│0.31│, ps≥0.325). Thus, while there might have been slight clearance of Aβ, but not tau. Neither Aβ nor tau load correlated with performance on the VM.

Halfway through the experiment, we developed a protocol that allowed us to assess 6e10 and glial cells on the same tissue, and thus collected glial data from that point forward but not prior, resulting in only half of the animals having glial stains done. Thus, we were underpowered for assessing both astrocytes and microglia. With this caveat in mind, we assessed the density of astrocytes measured by GFAP, which did not show a clear difference in any region measured (dCA1, vCA1, PPC, **Fig. 6D**, **Table 3**). Similarly, microglia measured by Iba1 did not show a clear difference in any region measured (**Fig. 6L**). Due to no clear differences and low n, we are not reporting statistics on glial cells as they are likely uninformative.

**Table 3.** Glial cell density averages. Average ± SEM glial cell density for GFAP Left hemisphere, GFAP right hemisphere, and Iba1 for each region measured.

|  | dCA1 | vCA1 | PC |
| --- | --- | --- | --- |
| GFAPL | 40 Hz: $1.4 \times 10^{-3} \pm 1.4 \times 10^{-4}$<br>sham: $1.1 \times 10^{-3} \pm 1.3 \times 10^{-4}$ | 40 Hz: $6 \times 10^{-4} \pm 6 \times 10^{-5}$<br>sham: $1.2 \times 10^{-3} \pm 1.1 \times 10^{-4}$ | 40 Hz: $1.2 \times 10^{-3} \pm 2.4 \times 10^{-4}$<br>sham: $1.1 \times 10^{-3} \pm 1.2 \times 10^{-4}$ |
| GFAPR | 40 Hz: $7 \times 10^{-4} \pm 1.4 \times 10^{-4}$<br>sham: $1.1 \times 10^{-3} \pm 8 \times 10^{-5}$ | 40 Hz: $6 \times 10^{-4} \pm 3 \times 10^{-5}$<br>sham: $1.5 \times 10^{-3} \pm 2.8 \times 10^{-4}$ | 40 Hz: $8 \times 10^{-4} \pm 1.0 \times 10^{-4}$<br>sham: $4 \times 10^{-4} \pm 5 \times 10^{-5}$ |
| Iba1 | 40 Hz: $7 \times 10^{-4} \pm 8 \times 10^{-5}$<br>sham: $7 \times 10^{-4} \pm 5 \times 10^{-5}$ | 40 Hz: $8 \times 10^{-4} \pm 1.2 \times 10^{-4}$<br>sham: $9 \times 10^{-4} \pm 1.4 \times 10^{-4}$ | 40 Hz: $1.3 \times 10^{-3} \pm 2.8 \times 10^{-4}$<br>sham: $8 \times 10^{-4} \pm 8 \times 10^{-5}$ |

## Discussion

Previously we found impaired HPC-PC interactions and decoupling of these events from behavior in 3xTg-AD mice and hints that HPC may be key in producing these deficits. Thus, we used 40 Hz optogenetic modulation in the HPC, achieved via stimulation to PV+ interneurons, to test the hypothesis that HPC targeted stimulation would be sufficient to rescue dysfunctional HPC-cortical coordination during sleep in female 6-month 3xTg-AD mice (Cushing et al., 2020). In support of our hypothesis, we found that HPC 40 Hz stimulated mice performed better on the VM than sham stimulated mice. Further, 40 Hz stimulated mice had intact HPC-PC interactions that appeared to be identical to NonTg animals from our previous work (Cushing et al., 2020). HPC-PC interactions were impaired in sham stimulated mice, similar to 3xTg-AD animals from our previous work (Cushing et al., 2020). However, while the HPC-PC interactions in NonTg mice correlated with improved performance on the task the subsequent day, in 40 Hz stimulated mice, this relationship between HPC-PC interactions and subsequent performance was absent. This suggests that after behavior becomes decoupled from HPC-PC interactions, potentially as a consequence of pathophysiology that occurred prior to the onset of 40 Hz stimulation, rescuing those interactions with 40 Hz stimulation does not restore the relationship with behavior. Finally, in 40 Hz stimulated mice, individual markers of memory reactivation in both HPC (SWR number) and PC (DWT number) now predict performance on the task the subsequent day. Together this pattern of results suggests that even very low levels of pathology may alter brain dynamics and that subsequent 40 Hz stimulation enables restored cognition in this altered system. Thus, in order to successfully treat AD, further mechanistic research at the systems level paired with cognitive assessment is of critical importance for improving the therapeutic potential of 40 Hz stimulation techniques and potentially also other therapeutic approaches.

We found that HPC-PC interactions during sleep, measured by SWR-DWT correlations, were rescued by 40 Hz modulation in CA1. However, unlike in NonTg mice (Cushing et al., 2020), this measure did not correlate with performance on the task the subsequent day. Similar to unstimulated 3xTg-AD mice, both 40 Hz and sham stimulated 3xTg-AD/ PV^cre^ mice showed no relationship between SWR-DWT correlations and performance on the task the subsequent day. Thus, pathology accumulation prior to the onset of 40 Hz stimulation may have produced changes in brain dynamics during sleep which persisted to some extent after 40 Hz stimulation was initiated leading to increased dependence on individual events for memory consolidation.

We found that the number of SWRs and DWTs correlated with improved performance on the VM in 40 Hz stimulated, but not sham stimulation mice. SWRs during both sleep and wakefulness have been shown to be positively correlated with spatial memory improvements (Jadhav et al., 2012; Buzsáki, 2015; Roux et al., 2017). Furthermore, optogenetically prolonging SWRs has resulted in improved performance on a memory task, while shortening or suppressing SWRs resulted in decreased performance (Ramadan et al., 2009; Ego-Stengel and Wilson, 2010; Fernández-Ruiz et al., 2019a). 40 Hz stimulation may result in an improvement of some aspect of the SWR other than number or rate, such as an improvement in power or temporal structure, which could result in the correlation with memory performance.

It has been suggested that memory formation in part involves coordination between slow gamma and SWRs in HPC, which may contribute to coordination with cortical activity (Carr et al., 2012; Pedrosa et al., 2022). Furthermore, slow gamma during SWRs has been shown to be impaired in rodent models of amyloid, APOE aggregation, and amyloid and tau aggregation, suggesting this dyscoordination may underlie impaired memory in AD (Gillespie et al., 2016; Iaccarino et al., 2016; van den Berg et al., 2023). Given that SWR-DWT temporal correlations no longer predicted improved behavioral performance in 40 Hz stimulated mice while total number of SWRs did, we could hypothesize that slow gamma during the SWR might have been affected by 40 Hz stimulation. However, recent work exploring the mechanism of slow gamma and SWR generation in the HPC indicates that gamma in HPC cannot occur during SWRs, given that opposite levels of acetylcholine seem to be critical for SWR and gamma generation, and that stimulating acetylcholine increases gamma and almost entirely abolishes SWRs (Gulyás et al., 2010; Schlingloff et al., 2014; Zhang et al., 2021). Thus, the 40 Hz changes during SWRs previously observed may reflect aspects of the SWR itself, and not a change in gamma. Thus, we did not assess gamma during SWRs. However, it is possible that 40 Hz treatment rescues other deficits in SWRs, such as power and temporal structures of SWRs further contributing to the shift in 40 Hz animals to correlations between individual events in HPC or PC and cognition (Witton et al., 2016).

Delta waves have similarly been correlated with memory performance in animals and humans (Ruch et al., 2012; Cushing et al., 2020). Furthermore, optogenetically stimulating delta waves during sleep in rats has also been shown to improve memory (Todorova and Zugaro, 2019). The finding that 40 Hz stimulation resulted in a correlation between DWTs and VM performance is unexpected given the 40 Hz stimulation was targeted to HPC. This suggests that 40 Hz stimulation of CA1 results in some change in DWTs in PC despite being several synaptic connections away. Consistent with this possibility, 40 Hz power was elevated in PC in 40 Hz HPC stimulated mice versus control mice.

However, other studies have noted that delta waves contribute to forgetting potentially unimportant information, and slow oscillations contribute to memory formation (Kim et al., 2019). These seemingly contradictory findings may be the result of some studies (including ours) not separating the detection of delta waves vs slow oscillations, since they have overlapping frequency ranges but have different wave forms (Dang-Vu et al., 2008; Kim et al., 2019). 40 Hz manipulation may be shifting towards slow oscillations and away from delta waves, leading to an increased proportion of memory consolidating reactivation events.

We found that spindle coupling with both DWTs and SWRs were similar across stimulation, and were not altered by 40 Hz stimulation. This is likely a result of spindle generation occurring in the thalamus, synaptically upstream of dCA1 where stimulation occurred (Tao et al., 2021). Given that HPC targeted 40 Hz modulation may not produce modulation in thalamus, this finding is not unexpected.

We found that 40 Hz stimulation of PV+ interneurons in CA1 did not have an impact on the number or rate of SWRs in CA1, or DWTs or spindles in PC. AD rodent models show a reduction in the number, frequency, and power of SWR (Nicole et al., 2016; Witton et al., 2016; Jones et al., 2019a; Caccavano et al., 2020). Additionally, some AD rodent models show a reduction in gamma power during SWRs (Gillespie et al., 2016; Stoiljkovic et al., 2018). 40 Hz stimulation effects on SWRs has not previously been reported, and our results indicate that 40 Hz optogenetic stimulation may not rescue SWR abundance. Sleep spindles also display abnormalities in both AD patients and AD rodent models (Gorgoni et al., 2016; Cushing et al., 2020; Liu et al., 2020) for review see (Weng et al., 2020). Some rodent models of AD also show reductions in delta power during sleep (Leparulo et al., 2019; Chen et al., 2023). 40 Hz stimulation effects on DWTs and spindles has not previously been measured, and our results indicate that 40 Hz stimulation of CA1 also may not have a direct effect on the number or rate of these markers of memory related population activity.

We found that while 40 Hz HPC stimulation had no effect on total sleep time, or proportion of sleep spent in SWS. However, it is possible that 40 Hz has some impact these measures because sleep time and time in SWS were only correlated with subsequent performance on our spatial navigation task in the 40 Hz stimulated mice and not sham mice. Previous studies, including our own, have found that AD has significant impact on sleep in both humans and animal models (Vitiello et al., 1990; Grace et al., 2000; Bonanni et al., 2005; Wisor et al., 2005; Jyoti et al., 2010; Roh et al., 2012; Rothman and Mattson, 2012; Lim et al., 2013; Rothman et al., 2013; Di Meco et al., 2014; Ju et al., 2014; Peter-Derex et al., 2015; Sethi et al., 2015; Varga et al., 2016; Holth et al., 2017; Mander et al., 2017; Kent et al., 2018; Olsson et al., 2018; Van Erum et al., 2019; Winsky-Sommerer et al., 2019; Wang and Holtzman, 2020; Kent et al., 2021). While other studies in humans have indicated that multi-sensory 40 Hz stimulation may impact sleep, we found no such results using optogenetic stimulation in our mice (Cimenser et al., 2020). This could be the result of our stimulation being targeted to the HPC, or could be related to the timing of our manipulation (when pathology levels were low). Related to the later point, 3xTg-AD mice actually sleep more than NonTg mice, similar to humans with white matter hyperintensities and free-water fractions, markers of cerebrovascular injuries in AD (Baril et al., 2024). Further, we did find that SWS as well as total sleep time did predict improved performance on the VM in 40 Hz stimulated mice, which does suggest that some aspect of sleep was improved and is consistent with a vast body of literature indicating sleep and specifically SWS improves cognitive performance and memory retention (Frankland and Bontempi, 2005; Diekelmann and Born, 2010; Westerberg et al., 2010; Ruch et al., 2012; Ju et al., 2013; Di Meco et al., 2014; Peter-Derex et al., 2015; Helfrich et al., 2018; Klinzing et al., 2019; Van Erum et al., 2019; Spanò et al., 2020; Chen et al., 2023). However, given that SWS and sleep time did not correlate in NonTg or 3xTg-AD mice previously, it is possible that this relationship between sleep and cognition is actually reflecting the relationship between memory related activity marker number (SWR and DWT) and performance. 3xTg-AD/ PV^cre^ mice that underwent 40 Hz optogenetic stimulation showed a decrease in Aβ load in vCA1 and PC compared to sham mice. Previous findings report that optogenetic stimulation of dCA1 with 40 Hz resulted in decreased Aβ load in dCA1 compared to random stimulation (Iaccarino et al., 2016). Our results do highlight that this clearance may spread further across the HPC-PC circuit than just dCA1. While we did not show clearance in dCA1, it is possible this is a result of the substantial amount of damage done to dCA1 when removing the recording array. However, some more recent studies have shown visual 40 Hz visual stimulation does not result in clearance of Aβ in any region (Soula et al., 2023; Yang and Lai, 2023), a failure to replicate the weaker clearance effects of visual 40 Hz stimulation reported previously (Iaccarino et al., 2016). It is possible that these failures to replicate are confined to visual only 40 Hz stimulation as 40 Hz stimulation with only one sensory stimulation, either visual or auditory, results in substantially less Aβ clearance (Martorell et al., 2019). Finally, our 40 Hz stimulation was delivered contralaterally, and thus would likely be weaker than the ipsilateral stimulation in (Iaccarino et al., 2016). Though it is also possible that our findings of reduced Aβ in some regions are an extension of these failures to replicate with visual only stimulation. Furthermore, Aβ load in dCA1, vCA1, and PC did not correlate with performance on the virtual maze. This is consistent with a great deal of work in both humans and animals (including our own data) showing the Aβ load does not correlate with or predict memory impairments (Terry et al., 1991; Gold et al., 2000; Gold et al., 2001; Giannakopoulos et al., 2003; Lin et al., 2009; Mormino et al., 2012; Huijbers et al., 2014; Stimmell et al., 2021), though there are some exceptions, including with our own data (Desikan et al., 2012; Cushing et al., 2020; Lussier et al., 2020).

Additionally, we found no reduction in tau load in any region measured. Visual 40 Hz stimulation has been previously reported to reduce tau phosphorylated at S202 and S400/T403/S404, but not S396 (Iaccarino et al., 2016). Our tau antibody binds tau phosphorylated at S202 and T205. It’s possible that the lack of reduction in tau load in our experiment is due to differences in stimulation (optogenetic vs visual) or differences in phosphorylation of tau (S202/T205 vs S202). Further, tau load did not correlate with performance on the VM. This is surprising, given that tau often correlates with memory performance in both mouse models (Arendash et al., 2004; Santacruz et al., 2005; Huber et al., 2018; Stimmell et al., 2021) and human patients (Brier et al., 2016; Hanseeuw et al., 2019; Sperling et al., 2019) for review see (Giacobini and Gold, 2013). Our previous work also suggests that the pattern of tau staining across a brain network may be a better predictor than tau accumulation in a single region (Stimmell et al., 2021). Similarly, there are exceptions in both AD patients and mouse models with absent correlations between tau load and behavioral impairments. It has been suggested that Aβ and tau have impacts on brain functions (e.g., sleep or synaptic plasticity) that indirectly lead to observed impairments and thus the most robust relationship is between cognition and the brain functional measures (Mander et al., 2015; Mander et al., 2016; Bejanin et al., 2017; Tiepolt et al., 2019; Winer et al., 2019; Lussier et al., 2020; Jagust et al., 2023). Combined with these findings, our results that neither Aβ nor tau directly correlate with memory performance, but aspects of memory reactivation do, is consistent with the work suggesting that other factors in disease may sometimes have greater impacts on cognition than accumulation of Aβ or tau.

In summary, we stimulated dorsal CA1 of HPC with 40 Hz or sham stimulation to attempt to rescue previously identified HPC-cortical interactions during sleep. We rescued both the hippocampal-cortical interactions, and additionally rescued performance on the spatial navigation task. However, performance on the task had become decoupled from HPC-cortical interactions and instead correlated with individual markers of memory reactivation. Thus, 40 Hz stimulation may have direct impacts on sleep related memory consolidation that may improve memory performance, though it does not necessarily restore previously abolished relationships between brain dynamics and memory. Our findings suggest that the timing of treatment interventions in AD may be of utmost importance and research of systems level mechanisms underlying AD deficits is critical for improving the therapeutic potential of 40 Hz and other AD treatments.

## Lead contact

Sarah Cushing

## Acknowledgements

Acknowledgements:

This research was supported by grants from NIA R00 AG049090, and R01 AG070094, and Florida DOH 20A09 to AAW, and NIA F31AG079619-01 to SDC. We thank Dr. Anabelle Singer for helpful comments on this manuscript.

